# A Bayesian framework for ranking genes based on their statistical evidence for differential expression

**DOI:** 10.1101/2025.01.20.633909

**Authors:** Franziska Hoerbst, Gurpinder Singh Sidhu, Thelonious Omori, Melissa Tomkins, Richard J. Morris

## Abstract

Advances in sequencing technologies have revolutionised our ability to capture the complete RNA profile in tissue samples, allowing for comparative analyses of RNA levels between developmental processes, environmental responses, or treatments. However, quantifying changes in gene expression remains challenging, given inherent biological variability and technological limitations. To address this, we introduce a Bayesian framework for differential gene expression (DGE) analysis. Our framework unifies and streamlines a complex analysis, typically involving parameter estimations and multiple statistical tests, into a concise mathematical equation. This allows statistical evidence for differential expression to be computed rapidly and transparently. We show how this approach can be used to evaluate variabilty of individual genes between replicates. A comparison of our framework with existing tools revealed differences that can be explained by commonly employed thresholds in other packages. This motivated us to explore ranking genes based on their statistical evidence as opposed to a binary classification as DEGs. Our analysis leads us to advocate the use of Bayes factors within a rank-based approach. This framework offers enhanced computational efficiency and delivers a transparent way to analyse, interpret and communicate DGE results.

## 1 Background

A common approach to unravel how a system works is to perturb it and measure the response. In molecular biology, a key response of interest is the change in gene regulation. RNA sequencing (RNA-Seq) is a powerful technique that privides in-sights into gene expression and transcriptional activity at a tissue sample level. By sequencing all RNA molecules present in a sample, RNA-Seq can reveal which genes are expressed or which genomic regions are transcribed at a specific time during development or in response to environmental stimuli [1]–[3]. RNA-Seq datasets can be compared to identify statistically significant changes in the amounts of RNA between the samples (differential gene expression, DGE), or over-representation analyses to identify interesting candidate genes for further investigations [4].

Excellent review articles are available that summarise RNA-Seq technologies, associated data analysis tools, and their successes and limitations [1], [5]–[14]. Given the complexity of the datasets, biological variation and inherent technical noise, it is hardly surprising that, despite sophisticated statistical approaches, reproducibility issues have been reported [4], [15]–[17]. Here, we examine differential gene expression analysis from a Bayesian perspective, Figure 1. We find that the question of whether two genes are differentially expressed can readily be cast as a binomial problem, leading to an analytical solution. We validated this new method using both simulated (Figure 2) and real data, and compared it to popular, existing methods (Figure 3). Although we find disagreements between the methods, we show that these differences arise due to choices of thresholds. We, therefore, advocate using Bayes factors as a means for ranking differentially expressed genes, making the usual log_2_ fold change and p-value cut-offs redundant.

**Figure 1:**
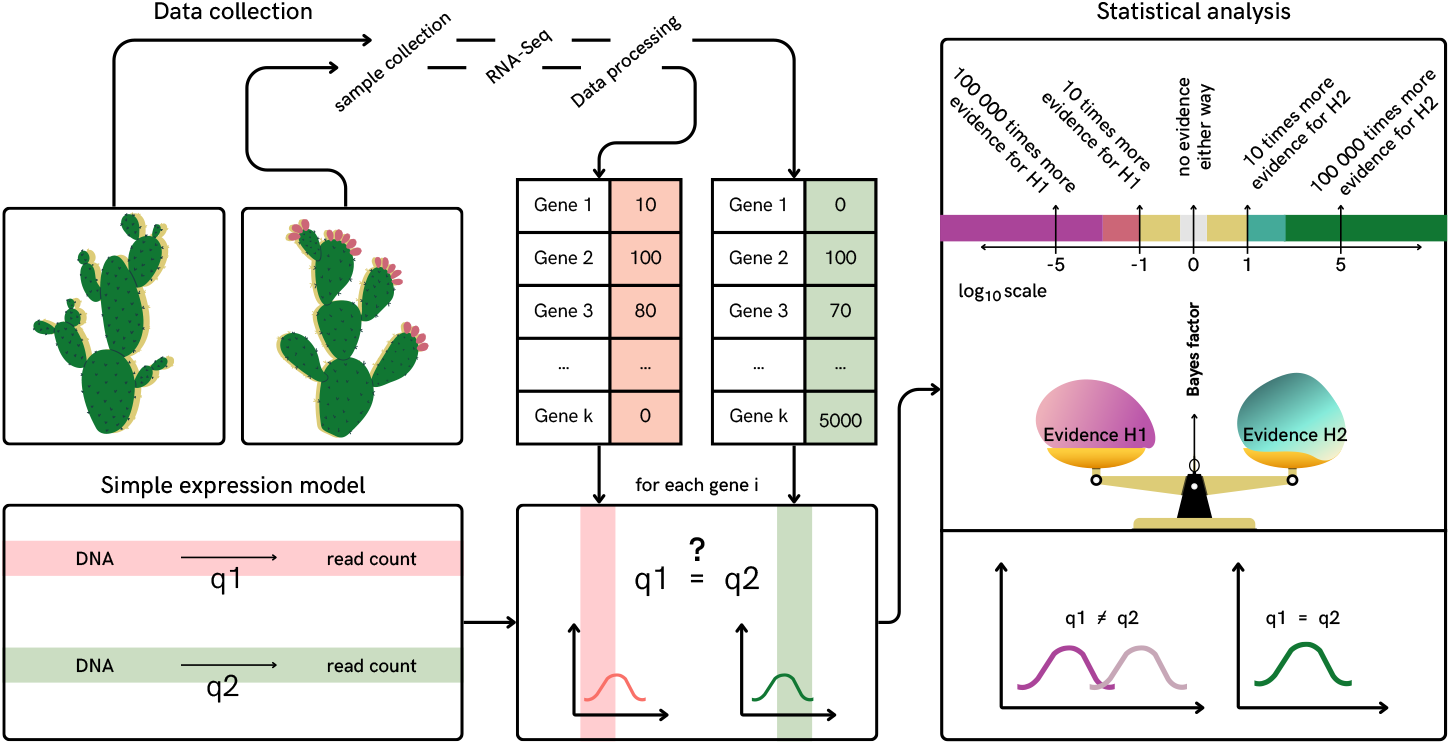
A schematic overview of differential gene expression analysis based on Bayesian model comparison. The analysis is built on a simple gene expression model for which we infer parameters from processed RNA-Seq data. We calculate the evidence (a measure for how compatible a model is with the data) for different models. The evidences for two alternative hypotheses and the corresponding models are compared by computing their ratio (Bayes factor). Hypothesis H1 assumes that the data can be explained by one gene expression parameter (*q*_1_ = *q*_2_), as opposed to hypothesis H2 for which two different gene expression parameters are needed to explain the data (*q*_1_ ≠ *q*_2_). H2 can be interpreted as a change in gene expression. Our knowledge of *q* is represented by a probability distribution over the parameter and not a single value. A negative log_10_ Bayes factor means that there is no change in expression, whereas a positive log_10_ Bayes factor marks a change. The more extreme the Bayes factor, the stronger the respective evidence. If log_10_ Bayes factors are around 0 there is no evidence either way. We can use Bayes factors to rank genes according to the confidence sin an expression change, and if wanted, for binary classifications.

**Figure 2:**
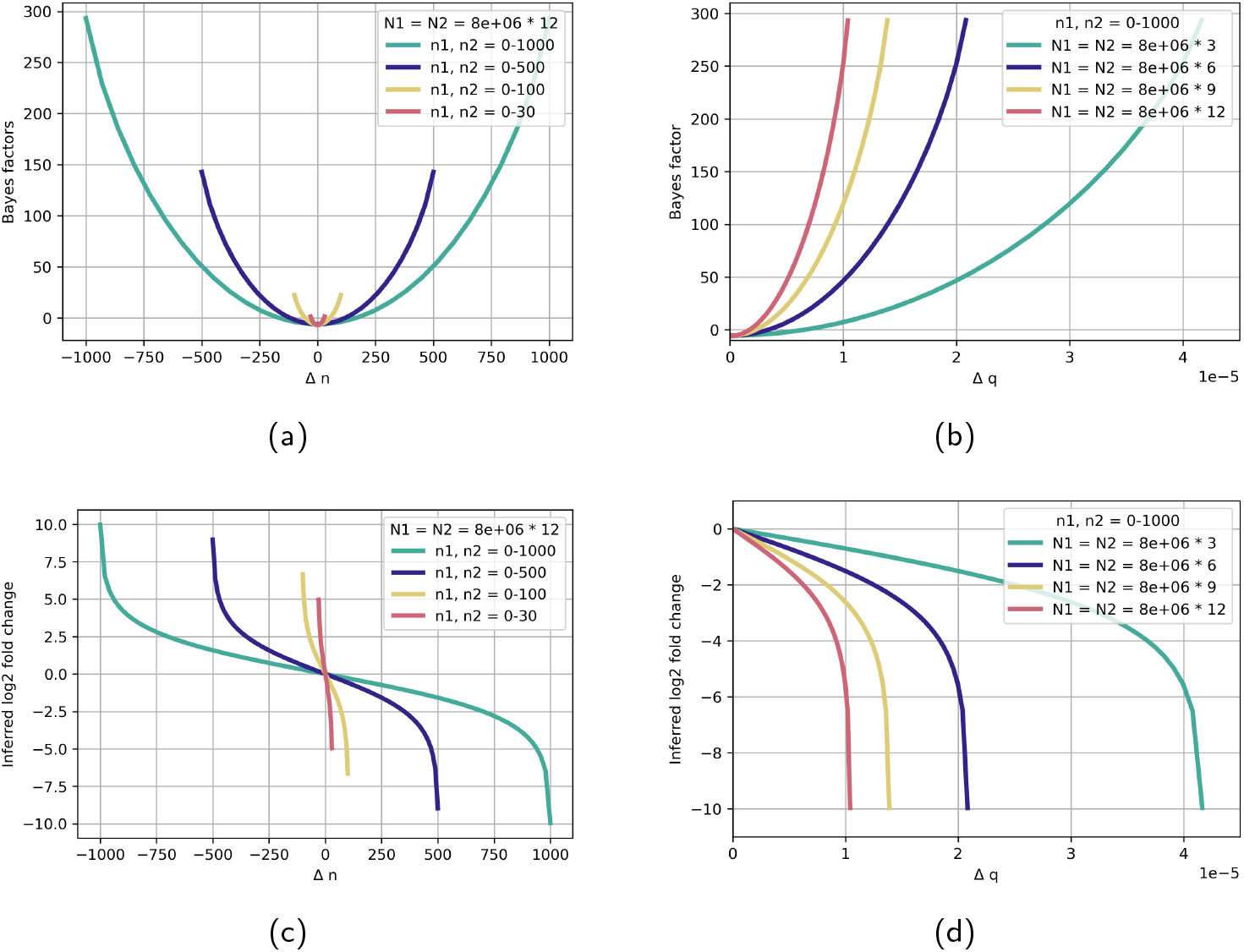
Bayes factors (log *BF*_21_) and absolute inferred fold change values increase with increased number of reads in RNA-Seq experiments. Bayes factors (log *BF*_21_) and inferred log_2_ fold change have been calculated following the presented equations for Bayes factors. In (a) and (c) the total number of reads in both *in silico* experiments is set to 8 × 10^6^ × 12. This number follows the average read depth in the RNA-Seq study of Schurch et al. [15] and their recommended number of 12 biological replicates.The four curves in each plot show the dependence of Bayes factor and log_2_ fold change on the the difference between the number of reads mapping to a gene in two different conditions, Δ*n* = *n*_2_ − *n*_1_. In (b) and (d) we show the behaviour of Bayes factors and inferred log2 fold change as a function of Δ*q* = (*n*_1_ + 1)*/*(*N*_1_ + 2) − (*n*_2_ + 1)*/*(*N*_2_ + 2) for different total read depths. We document what happens if the total read depth between conditions or treatments varies in Figures 6 and 7 in the Appendix.

**Figure 3:**
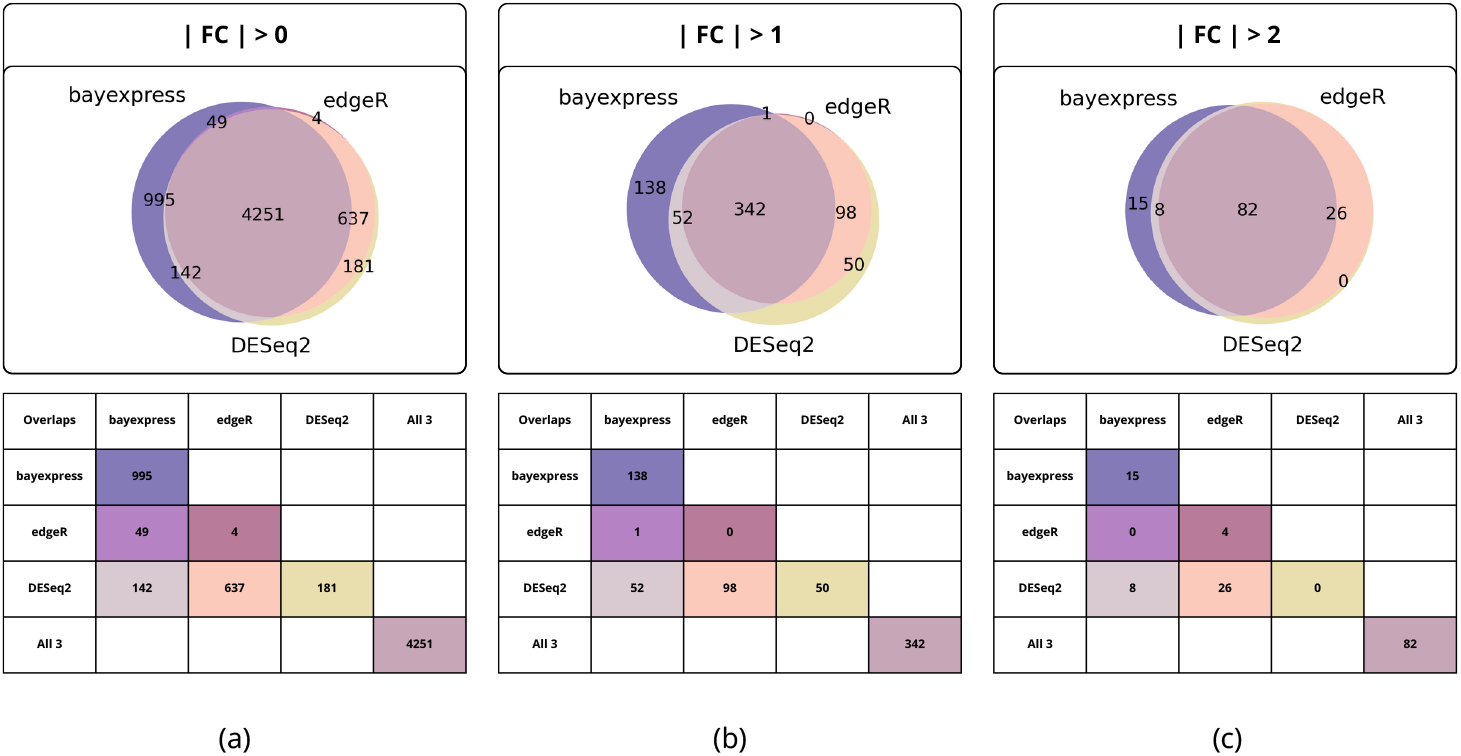
The popular tools *DESeq2* and *edgeR* and the presented Bayesian framework *bayexpress*, show disagreements in the identification of DEGs between WT and a SNF2-mutant in yeast. Interestingly, the discrepancies can be explained by small differences in log_2_ fold change values (see Supplementary Figure 8) that are amplified arbitrary classification thresholds. In the top panel, we show Venn diagrams of classified DEGs, in the bottom panel the same data is shown in table format. The three diagrams show three different sets of criteria for DEG: (a) Significant change (p-value *<* 0.05, or *BF*_21_ *>* 1) and | log_2_ fold change | *>* 0, (b) Significant change (p-value *<* 0.05, or *BF*_21_ *>* 1) and | log_2_ fold change | *>* 1, (c) Significant change (p-value *<* 0.05, or *BF*_21_ |*>* 1) and log_2_ fold change |*>* 2. In total there are 7126 yeast genes, the experiment had 42 WT replicates, and 44 SNF2-mutant replicates.

## 2 Results and Discussion

### 2.1 Differential gene expression analysis can be cast into the framework of Bayesian inference and model comparison

Consider two experiments with total RNA-Seq read counts equal to *N*_1_ and *N*_2_. We want to know whether gene *i* has statistically different read counts, *n*_*i*1_ and *n*_*i*2_, between the two experiments. The sum of all reads over all genes being the total number of reads, *N*_1_ = ∑_*i*_ *n*_*i*1_ and *N*_2_ = ∑_*i*_ *n*_*i*2_.

To describe experiments, we assign to every gene *i* an expression probability *q*_*i*_. This probability captures all events from transcription to data processing,

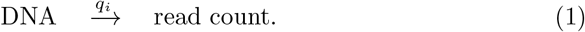

The expression probability can be different for every gene and change between experiments. As we show below, it is important to consider the total read counts, *N*_1_ and *N*_2_, as these numbers influence the expected change, e.g. 1 out of 100 has a different relative error to 100 out of 10,000 despite the ratio being the same.

For a known, fixed *q*_*i*_, we can describe the probability of *n*_*i*_ number of reads mapping to a gene *i*, out of total reads in a sample *N* with a binomial distribution,

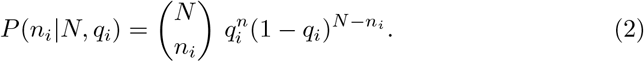

The underlying model assumes that a read maps to gene *i*, with probability *q*_*i*_, and does not, with probability 1 − *q*_*i*_. This implies that if we know the gene expression probability *q*_*i*_ of gene *i* and the total number of all reads *N* in an RNASeq experiment, we can compute the probability of any number of RNA-Seq reads *n*_*i*_ mapping to gene *i*.

Thus, in a differential gene expression experiment we want to answer the question of whether two datasets *D*_1_ (represented by *N*_1_ and for each gene *i* the read count *n*_*i*1_) and *D*_2_ (*N*_2_, *n*_*i*2_), arose from the same probability distribution *q*_*i*_,

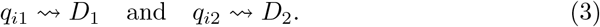

Therefore, our problem is to decide whether

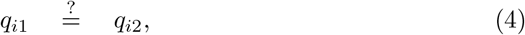

which is equivalent to asking whether the expression probability *q*_*i*_ of a gene changed between two datasets. We can calculate a Bayes factor for each gene, describing how much the RNA-Seq data supports one hypothesis over the other.

**Hypothesis H**_**1**_ states that the data from both experiments can be explained by one expression probability, *q*_*i*_, for each gene *i*,

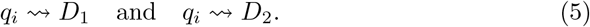

RNA-Seq data from the first and second experiment, i.e. mRNA levels of gene *i*, are statistically consistent, and we have no statistical support for a difference between them.

**Hypothesis H**_**2**_ states that the data are best explained by different expression probabilities for each experiment and each gene,

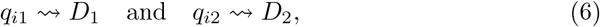

with *q*_*i*1 ≠_ *q*_*i*2_. In this case, the RNA-Seq data are unlikely to have arisen from just one *q*_*i*_, requiring a separate model for each dataset. Hence the data support a difference in gene expression between the samples.

The Bayes factor (*BF*_21_) is a ratio of the statistical evidences *P* (*D*_1_, *D*_2_|*H*_1_) and *P* (*D*_1_, *D*_2_|*H*_2_) supporting the two hypotheses. The full derivation can be found in the Appendix.

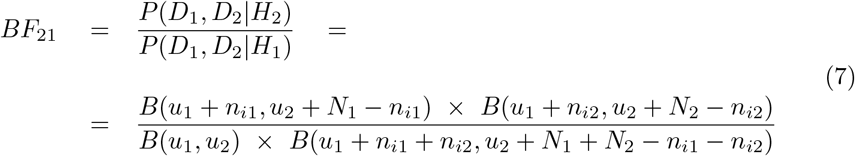

As is common practice in differential gene expression analysis, we proceed to calculate an inferred log_2_ fold change (*iFC*),

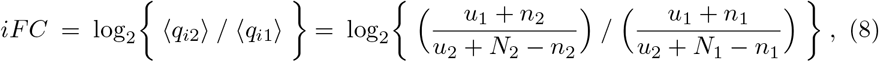

where *u*_1_ and *u*_2_ are hyperparameters (set to 1), see Appendix. Note that in contrast to other packages, we infer the fold change directly from read counts using Laplace’s rule of succession (equivalent to Bayesian inference using a uniform prior).

This approach allows for replicates to be handled by summing the *N* and *n*_*i*_ values. From these combined values, we can infer an average expression rate, log_2_ fold change (*iFC*) and compute a Bayes factor (*BF*_21_) between different experiments.

### 2.2 Ranking genes according to statistical evidence for expression change

The Bayes factor, *BF*_21_ = *P* (*D*_1_, *D*_2_|*H*_2_)*/P* (*D*_1_, *D*_2_|*H*_1_), provides a measure of confidence in one hypothesis over another after seeing the data, when both hypotheses were equally probable *a priori*. We log_10_ transform the Bayes factors. A log *BF*_21_ of 0 means there is an equal probability for both hypotheses, whereas a log *BF*_21_ *>* 0 favours Hypothesis 2 (change in gene expression) and log *BF*_21_ *<* 0 favours Hypothesis 1 (no change in gene expression), see Figure 1. Assigning each gene a log_10_ Bayes factor (log *BF*_21_) enables ranking them according to the evidence supporting gene expression change given the RNA-Seq data.

Should a simple ‘change’ or ‘no change’ outcome be preferred, Bayes factors can be used as criteria for a binary classification of DEGs. Where necessary in this study, we followed published literature [18], [19] and chose a cut-off of log *BF >* 1 → differentially expressed gene (DEG).

### 2.3 Ranking genes according to their variability across replicates

Biological and technical noise can introduce variation between replicates that can lead to potential false positives in DGE analysis [4], [15]–[17]. The presented framework allows for quantification of this variability. Using the same equations as derived above, we can ask whether two replicates have consistent expression for each gene. The framework can be extended to ask whether any number of replicates, *k*, have consistent expression.

We define two hypotheses. **Hypothesis H**_**1**_ states that the data from all replicates can be explained by a statistical model with only one expression probability *q* for each gene *i*,

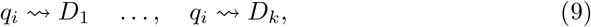

i.e the data are consistent with the biological and technical variance that might be expected between replicates. **Hypothesis H**_**k**_ states that the data are best explained by a separate expression probability for each replicate,

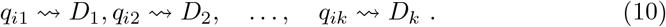

For each gene we calculate a Bayes factor, quantifying the evidence for Hypothesis *H*_*k*_ (*P* (*D*_1_, …, *D*_*k*_|*H*_*k*_)) over Hypothesis *H*_1_ (*P* (*D*_1_, …, *D*_*k*_|*H*_1_)), i.e. how consistent the expression is between replicates, from the RNA-Seq data,

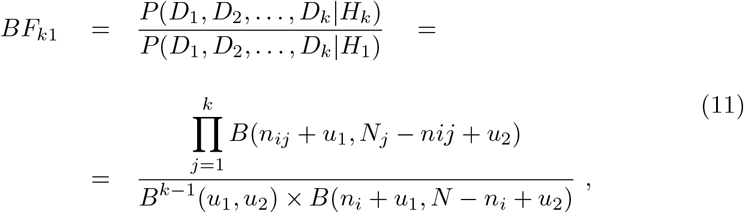

with *N* = ∑_*r*_ *N*_*r*_ and *n*_*i*_ = ∑_*r*_ *n*_*ir*_ for all replicates *r*. The derivation for this extension can be found in the Appendix.

In theory, the number of expression probabilities in a model can be between 1 and the number of replicates *k*.

The above formulism can in principle handle any arbitrary subsets of the data, however, as the number of subsets increases combinatorially with *k*, we limit ourselves to the extreme case of all replicates being the same (*H*_1_) or different (*H*_*k*_). If the replicates are consistent, we can sum all *n*_*ir*_ for replicate *r* for each gene *i*, and likewise *N* = *N*_*r*_, for calculating Bayes factors (*BF*_21_) for DGE analysis.

### 2.4 The more the data, the stronger the evidence

We investigated the general behaviour of Bayes factors and inferred log_2_ fold change and their relationships for different total read depths and number of reads mapping to single genes, see Figure 2. Furthermore, we investigated the impact of varying total read depth between conditions or treatments, see Figures 6 and 7. As expected, the evidences grow with available data and so do the absolute Bayes factors. Due to the chosen statistical frameworks and normalisation steps other statistical frameworks cannot account for effects of differing total read depths.

### 2.5 Rank-based approaches provide an alternative for communication of DGE results

Many tools are available for differential gene expression analysis [20]–[25]. Currently, *DESeq2* [17], [26] *and edgeR* [25], [27] *are two of the most popular software packages. Therefore, we compared Bayes factors for differential gene expression with these two approaches*.

*We conducted differential gene expression analysis on a yeast dataset collected by Schurch et al. [15]*. *We took the complete set of 42 wild-type (WT) and 44 mutant replicates and analysed the data using DESeq2, edgeR*, and the presented Bayesian framework that we call *bayexpress*. First, we compared the similarities in ranking sgenes by Bayes factors, p-value and log_2_ fold change values using a rank comparison algorithm [28], [29]. We find that ranking by log_2_ fold change leads to similar results for all three approaches, Supplementary table 1. However, Bayes factors show no similarity with other metrics, Supplementary table 1, suggesting they are capturing something distinct from fold change and p-values alone. Interestingly, we also observe no similarity in the rankings by p-values of *DESeq2* and *edgeR*.

**Table 1:** We have calculated rank-biased overlaps (RBO) [28], [29] between all metrics (columns on the outside) used in our comparison (p-values, Bayes factors, fold change values). The table is ordered according to RBO values (0 less ranking similarity1 to 1 highest ranking similarity), which we have calculated for a range of the *p* parameter (top row) influencing the rank-weighting. Ranking by fold change values shows the highest similarities between the 3 methods examined. We can also see how ranking by Bayes factors provides an alternate ranking, incorporating more information than just fold change.

| Rank 1 | p = 0.1 | p = 0.2 | p = 0.3 | p = 0.4 | p = 0.5 | p = 0.6 | p = 0.7 | p = 0.8 | p = 0.9 | Rank 2 |
| --- | --- | --- | --- | --- | --- | --- | --- | --- | --- | --- |
| p_DESeq2 | 3.86949e-27 | 1.33586e-19 | 3.56137e-15 | 5.13737e-12 | 1.51840e-09 | 1.66178e-07 | 0.000009 | 0.000316 | 0.007996 | BF_21 |
| p_DESeq2 | 4.18664e-25 | 3.58975e-18 | 4.18616e-14 | 3.30502e-11 | 6.01834e-09 | 4.37710e-07 | 0.000017 | 0.000450 | 0.009873 | iFC |
| p_DESeq2 | 4.22498e-25 | 3.71855e-18 | 4.51485e-14 | 3.75274e-11 | 7.25692e-09 | 5.64479e-07 | 0.000024 | 0.000666 | 0.014616 | FC_DESeq2 |
| p_DESeq2 | 8.37325e-25 | 7.17876e-18 | 8.36678e-14 | 6.59488e-11 | 1.19684e-08 | 8.65255e-07 | 0.000034 | 0.000854 | 0.016997 | FC_edgeR |
| p_DESeq2 | 7.75276e-15 | 6.83276e-11 | 1.41918e-08 | 6.33005e-07 | 1.21337e-05 | 1.36473e-04 | 0.001070 | 0.006639 | 0.039103 | p_edgeR |
| BF_21 | 1.64092e-06 | 5.15387e-05 | 3.82558e-04 | 1.56736e-03 | 4.62243e-03 | 1.10762e-02 | 0.023282 | 0.046644 | 0.103049 | iFC |
| FC_DESeq2 | 1.64092e-06 | 5.15477e-05 | 3.83066e-04 | 1.57602e-03 | 4.69828e-03 | 1.15089e-02 | 0.025103 | 0.052900 | 0.124904 | BF_21 |
| FC_edgeR | 1.64092e-06 | 5.15478e-05 | 3.83079e-04 | 1.57657e-03 | 4.70814e-03 | 1.16076e-02 | 0.025758 | 0.056049 | 0.136307 | BF_21 |
| p_edgeR | 1.65327e-06 | 5.31070e-05 | 4.09344e-04 | 1.77062e-03 | 5.61927e-03 | 1.47878e-02 | 0.034520 | 0.074156 | 0.148293 | BF_21 |
| p_edgeR | 3.26607e-03 | 1.29437e-02 | 2.92657e-02 | 5.31775e-02 | 8.65034e-02 | 1.31858e-01 | 0.191698 | 0.264505 | 0.326795 | iFC |
| FC_DESeq2 | 3.26620e-03 | 1.29510e-02 | 2.93392e-02 | 5.35372e-02 | 8.76936e-02 | 1.34932e-01 | 0.198327 | 0.276541 | 0.345219 | p_edgeR |
| p_edgeR | 3.26620e-03 | 1.29510e-02 | 2.93405e-02 | 5.35531e-02 | 8.77947e-02 | 1.35374e-01 | 0.199826 | 0.280713 | 0.355188 | FC_edgeR |
| FC_edgeR | 9.99980e-01 | 9.99692e-01 | 9.98481e-01 | 9.95328e-01 | 9.88896e-01 | 9.77466e-01 | 0.958747 | 0.929436 | 0.878415 | iFC |
| FC_DESeq2 | 9.99980e-01 | 9.99692e-01 | 9.98481e-01 | 9.95331e-01 | 9.88928e-01 | 9.77706e-01 | 0.960108 | 0.935577 | 0.901487 | iFC |
| FC_DESeq2 | 1.00000e+00 | 9.99997e-01 | 9.99992e-01 | 9.99928e-01 | 9.99565e-01 | 9.98083e-01 | 0.993253 | 0.980167 | 0.951003 | FC_edgeR |

If we want to compare the results of our ranking-based framework with traditional classification approaches we can follow published literature [18], [19] and choose a cut-off of log *BF >* 1 to mark significant changes in expression, analogous to p-value cut-offs for *edgeR* and *DESeq2* outputs. Additionally, we can set log_2_ fold change cut-offs in all three approaches. Figure 3 shows the overlaps of the classification results with varying fold change cut-offs. The agreement between *DESeq2, edgeR* and *bayexpress* overall is 68% for the three groups in Figure 3a, 50% in 3b, and 63% in 3c. However, we observed that *edgeR* and *DESeq2* had better overlaps: 93% for Figure 3a, 81% for 3b, and 93% for 3c. Whilst this is expected given the similarity of their underlying statistical models, the classification discrepancies with *bayexpress* were not clear and led us to investigate further.

In Figure 4 and 5 we show a selection of genes that are inconsistently labeled by the three approaches. For instance, the gene in Figure 4a scores a high Bayes factor for expression change between the 42 wild-type and 44 mutant replicates, with an inferred log_2_ fold change above 1, which is not identified by *edgeR* and *DESeq2* due to minor differences in log_2_ fold change values (see Figure 5 (d)). Similarly, 4 (c) shows a gene that is identified by *edgeR* and *DESeq2* but not the Bayesian framework, when classification thresholds are introduced. We observe that all differences in classifications are a result of variations in the estimated log_2_ fold change values between the approaches (see Supplementary Figure 8), and/or pre-filtering of the data (*edgeR* was set to filter genes with 0 values in any of the replicates). Both of these genes have roughly halved or doubled their expression but are classified differently. All these differences we identified in the results between approaches can be explained by the choice to classify and with that the need to apply classification thresholds.

**Figure 4:**
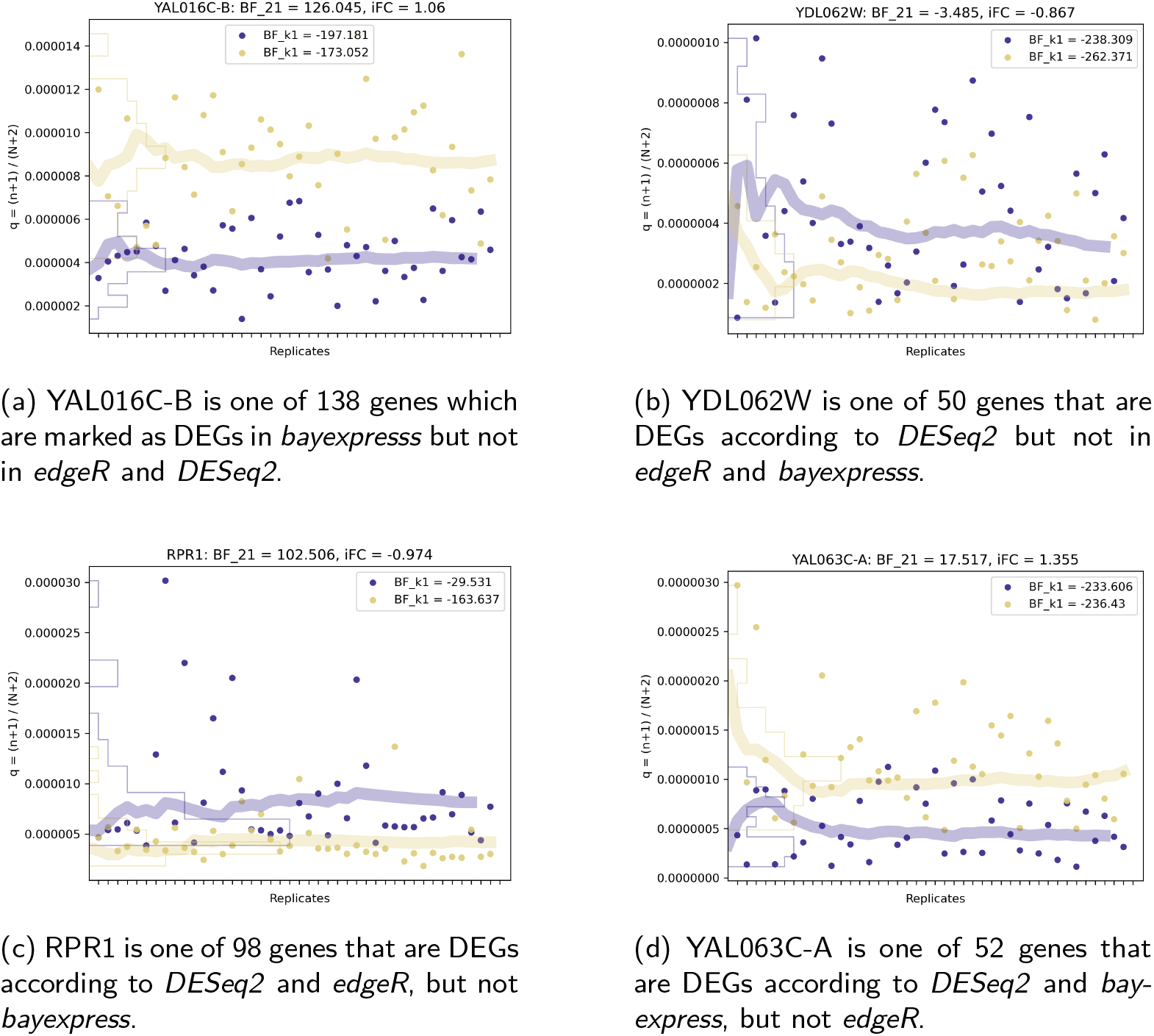
Disagreements in the classification of DEGs arise because of pre-filtering and small differences in fold change values. Here, and continued in Figure 5, we plot the inferred expression probabilities, *q*, for 7 example genes across all replicates (42 WT, 44 SNF2-mutant) in a yeast experiment [15]. The WT is shown in purple and the SNF2-mutant in yellow. Note the different scales on the y-axis. The fine lines are density histograms along the y-axis, and the thick lines are estimated means of *q*, updated with each new replicate. For each gene we show the Bayes factor for differential expression *BF*_21_ and an inferred log_2_ fold change at the top, both are calculated taking all 42/44 replicates into account. Bayes factors for consistency *BF*_*k*1_ of replicates can be found for each genotype in the boxes. The example genes have been selected to cover all sets in the Venn diagram in Figure 3b. Continued in Figure 5.

**Figure 5:**
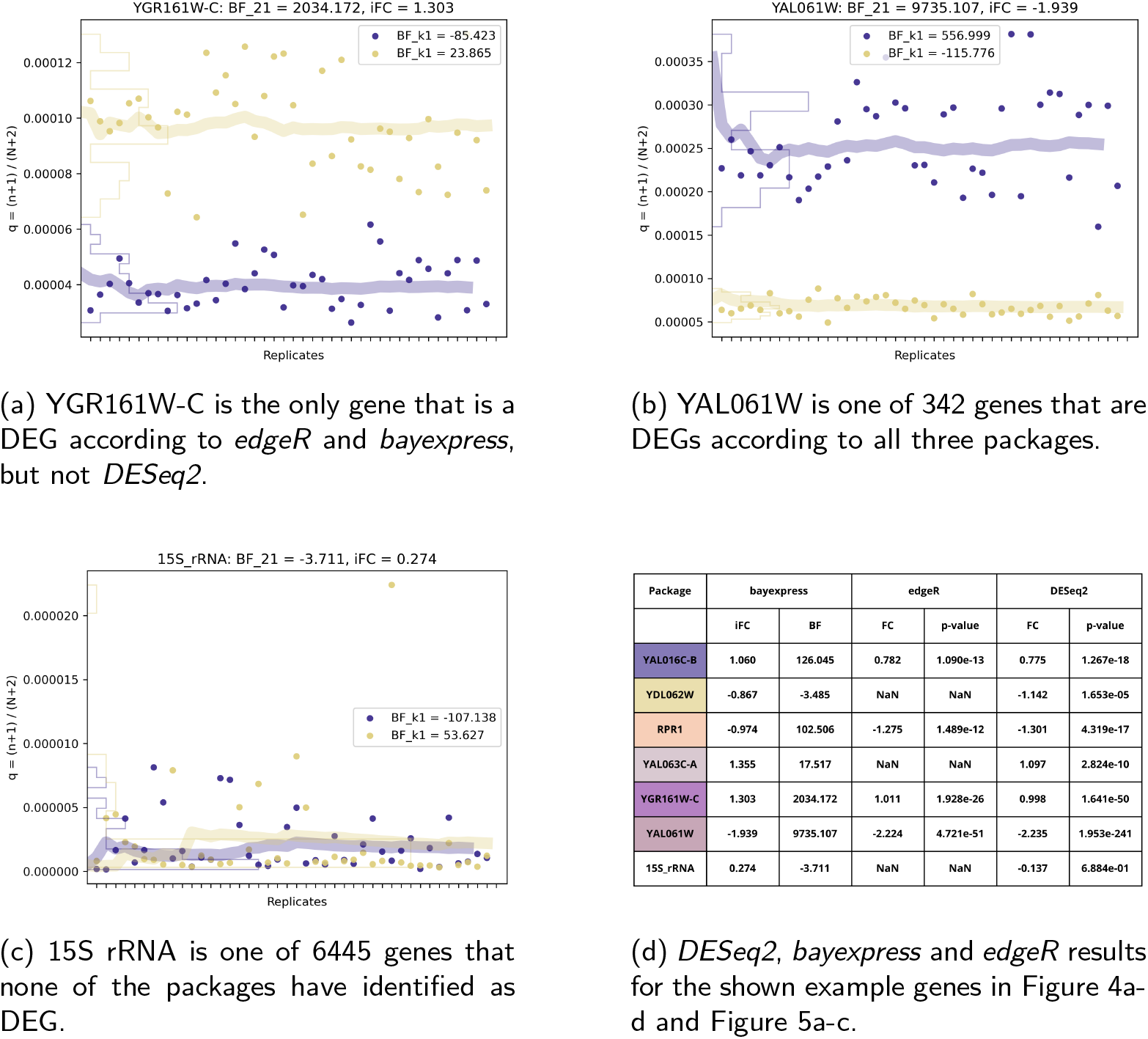
Disagreements in the classification of DEGs arise because of pre-filtering and small differences in fold change values. Here, as a continuation of Figure 4, we plot the expression probabilities, *q*, for 7 example genes across all replicates (42 WT, 44 SNF2-mutant) in a yeast experiment [15]. The WT is seen in purple, and the SNF2-mutant in yellow. Note the different scales on the y-axis. The fine lines are density histograms along the y-axis, and the thick lines are estimated means of *q*, updated with each new replicate. For each gene we show the Bayes factor for differential expression (*BF*_21_) and an inferred log_2_ fold change at the top, both are calculated taking all 42/44 replicates into account. Bayes factors for consistency of replicates (*BF*_*k*1_) can be found for each genotype in the boxes. The example genes have been selected to cover all sets in the Venn diagram in Figure 3b.

**Figure 6:**
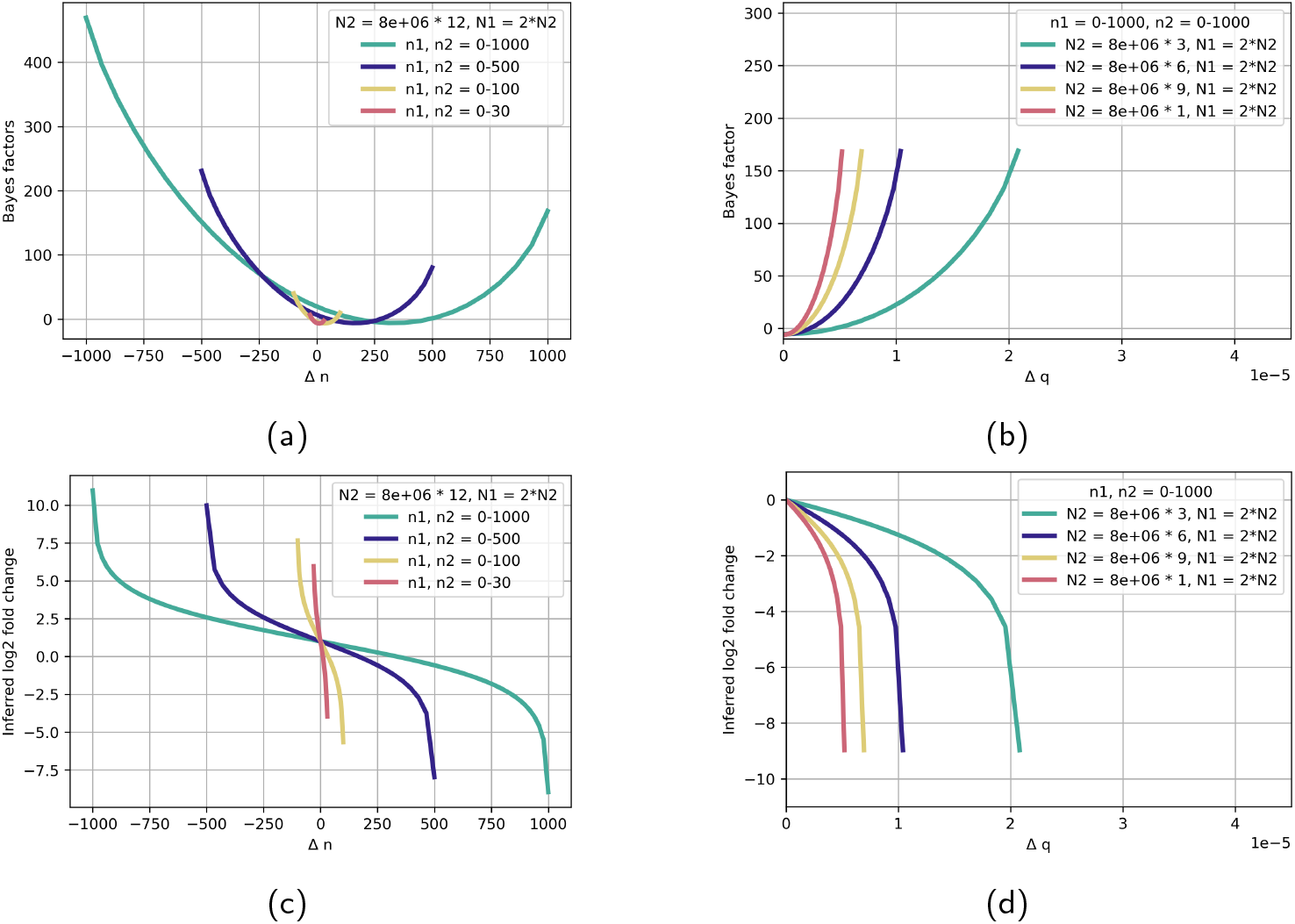
Differences in overall read depths between experiments (while *n*_1_ and *n*_2_ stay in similar magnitudes) are reflected in Bayes factors. In line with Figure 2, we see that Bayes factors and absolute inferred fold change values rise with more data (increased number of reads in RNA-Seq experiments). Here, one experiment has double the total number of reads compared to the other. The number of reads mapping to genes, *n*_1_ and *n*_2_, keep the same ratios. In (a) and (c) the total number of reads in the *in silico* experiments is set and we see variation in the read depths per genes. The four curves show Bayes factor and log_2_ fold change as a function of the difference between the number of reads mapping to a gene in two different conditions, Δ*n* = *n*_2_ − *n*_1_. In (b) and (d) we show Bayes factors and inferred log_2_ fold change as a function of the differences in gene expression probabilities *q*, Δ*q* = ((*n*_1_ + 1)*/*(*N*_1_ + 2)) − ((*n*_2_ + 1)*/*(*N*_2_ + 2)) for different total read depths, again *N*_1_ ≫ *N*_2_.

**Figure 7:**
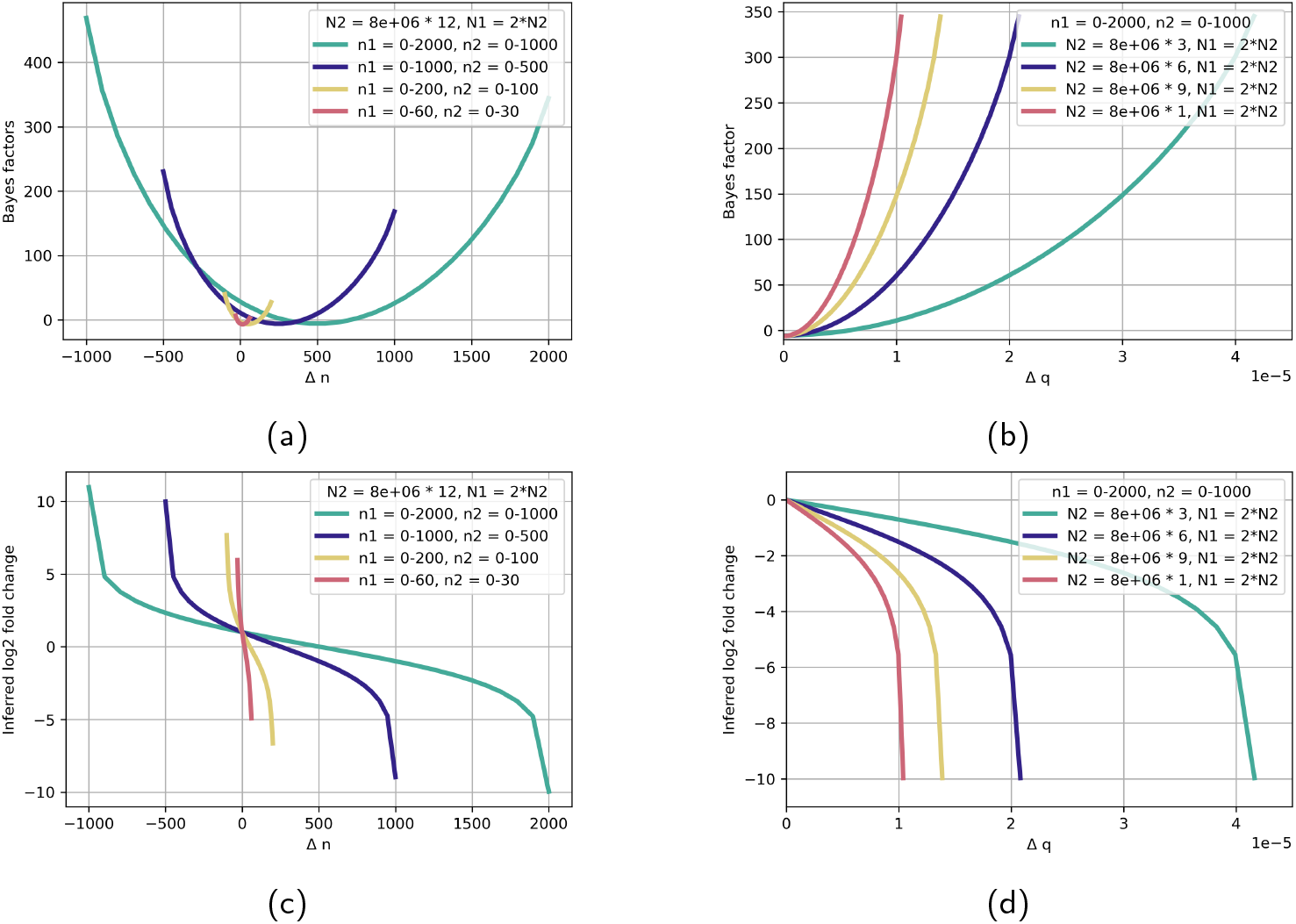
Differences in overall read depths in combination with *n*_1_ and *n*_2_ rising in proportion are reflected in Bayes factors. In line with Figure 2, we see how Bayes factors and absolute inferred fold change values rise with more data (increased number of reads in RNA-Seq experiments). Here, one experiment has double the total number of reads compared to the other. In contrast to Figure 6, here *n*_1_ and *n*_2_ are in relation to the raised read depth in one experiment. In (a) and (c) the total number of reads in the *in silico* experiments is set and we see variation in the read depths per genes. The four curves show Bayes factor and log_2_ fold change as a function of the difference between the number of reads mapping to a gene in two different conditions, Δ*n* = *n*_2_ − *n*_1_. In (b) and (d) we show Bayes factors and inferred log_2_ fold change as a function of Δ*q* = ((*n*_1_ + 1)*/*(*N*_1_ + 2)) − ((*n*_2_ + 1)*/*(*N*_2_ + 2)) for different total read depths, again s*N*_1_ ≫ *N*_2_ and *n*_1_ ≫ *n*_2_.

**Figure 8:**
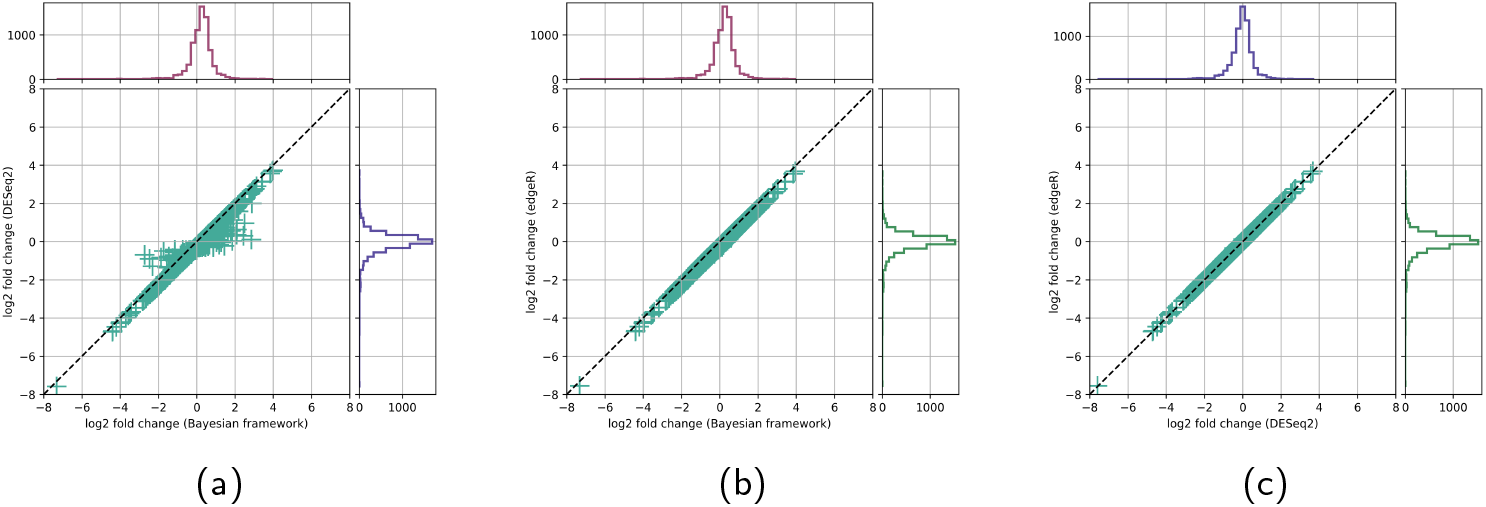
There are small differences in the values of the inferred log_2_ fold change of the presented Bayesian framework and log_2_ fold changes calculated by *DESeq2* and *edgeR*. These differences lead to disagreements in DGE classifications. The three log_2_ fold changes are shown against one another. Data: 7126 yeast genes (42 WT replicates, 44 SNF2-mutant replicates) [15]. (a) *DESeq2* vs. Bayesian framework. Overall, the fold change values are shifted and do not match perfectly. We checked the genes where *DESeq2* and the Bayesian framework do not agree (off-diagonal) and found they were genes with no reads in at least one replicate. For such genes, the effect of inferring the log_2_ fold change using Laplace’s rule of succession (which results from the choice of uniform prior) causes the largest difference. (b) *edgeR* vs. Bayesian framework. Again, we can see an overall shift between the fold change value calculated by *edgeR* and the inferred fold change. Filtering genes with zero reads in *edgeR* results in no genes with off-diagonal values. (c) The fold change values of *DESeq2* and *edgeR* match nearly perfectly.

Casting differential gene expression into a binary classification framework is an oversimplification of the insights we can obtain from RNA-Seq data and inevitably leads to loss of interpretable information for the user. To alleviate this disagreement in classification, we advocate ranking-based methods to communicate DGE results.

## 3 Materials and methods

All data (experimental and simulated) with the scripts to repeat the study and reproduce figures can be found on our GitHub.

### 3.1 How to use Bayes factors for differential gene expression analysis

The presented equations can be implemented in just a few lines of code and be used to calculate Bayes factors for differential gene expression, Bayes factors for consistency of replicates, and inferred log_2_ fold change values, for all genes from processed RNA-Seq data (read counts). We provide a Python and R implementation on our GitHub.

### 3.2 Yeast data

The package comparison was inspired by a yeast study conducted by Schurch et al. [15], [30] in 2016. They performed RNA-Seq on 42 wild type (WT) and 44 SNF2-mutant replicates and carried out a bootstrapping study to find out how many biological replicates RNA-Seq studies need and which packages to use for the analysis. Thanks to their detailed documentation and data availability, we could build on their developments in the presented study.

We used the fully processed data, downloaded from their GitHub. Originally they had 48 replicates but only worked with 42/44, as documented in [15], [30], which we followed.

## 4 Conclusions

In this paper we described an exact solution for the two-sample-test problem in differential gene expression analysis on RNA-Seq data. We tested it on both simulated (Figure 2) and real RNA-Seq data, and compared to existing packages (Figure 3). Our approach provides a framework for ranking genes based on the statistical support, characterised by Bayes factors, for change in gene expression.

The underlying model is a binomial distribution, which is, with no further knowledge, an optimal (maximum entropy, least-biased) probability assignment [18]. Other approaches have used a negative binomial to capture the overdispersion, relative to Poisson, of normalised expression values between biological replicates [17], [25], [26]. However, this is done without reference to the total read counts.

Variability in RNA-Seq experiments has been well-documented [4]–[8], [16], [17], [20], [21], [25], [26], yet how much of this variation is technical or biological remains under discussion [31]–[34]. This leads to challenges in the interpretation of the DGE results due to potential consistency issues. Bayes factors can be used to quantify the consistency of replicates and rank genes according to their variability. The quality of this ranking is dependent on available information about the variability of a gene, i.e. the number of replicates. This part of the framework could be extended to identify DEGs in multiple comparison studies.

Our analysis revealed divergent results between the three DEG approaches we investigated. Some of these can be attributed to how the data is pre-filtered (e.g. excluding genes with 0 reads). Small differences in the calculation of log_2_ fold changes also contribute to changes in the classification once cut-offs are introduced (Figure 8). Both of these issues are reflected in the examples in Figure 4 and 5. Introducing log_2_ fold change value cut-offs (e.g. | log_2_ fold change | *>* 1 or | log_2_ fold change | *>* 2) for binary classification often amplifies these discrepancies, even though genes that have not doubled or quadrupled their expression can have more supporting evidence for a change in gene expression. This introduces the possibility of loosing interesting candidates with good statistical evidence for change. The discussion about the accuracy of RNA-Seq below certain magnitudes might have motivated these thresholds as a control for false positives [35], [36], even though small changes can have significant biological effects. Higher numbers of replicates have been proposed to increase the accuracy and reproducibility [15], but for in a limited replicate scenario ranking genes by Bayes factors can help identify candidates without loosing interesting genes to cut-offs. Our analysis suggests that Bayes factors incorporate additional information, hence providing an alternative to ranking just base on fold change, see Supplementary table 1.

There is potential for further expansions of the presented framework. There are numerous articles discussing the analysis of single-cell RNA-Seq data [37]–[40]. While our framework is not specifically tailored to single-cell sequencing data, it holds potential for adaptation to such data. Furthermore, despite us not incorporating additional information in the prior (i.e. we chose a uniform prior), there is huge potential in doing so in the future. Information about the system (e.g. known expression changes between tissues or organisms) and experimental design (e.g. batch-effects or alike) can be accounted for in the analysis. If required, Bayes factors for differential gene expression (*BF*_21_) can be plotted against inferred log_2_ fold change values to mimic Volcano plots [13], [41]. Furthermore, the extended framework using *k* different models could be used to quantify differential gene expression in multiple comparison studies. Employing a ranking-based framework allows for comparison using rank comparison algorithms [28], [29] offering new avenues for interpretation of results in RNA-Seq studies.

## 5 Appendix

### 5.1 Deriving Bayes factors for differential gene expression

The expression probability, *q*_*i*_, can be inferred from RNA-Seq data using Bayes’ theorem. We use *θ*_*i*_ to denote the range of possible values of *q*_*i*_ (between 0 and 1) and use a probability distribution over *θ*_*i*_, *P* (*θ*_*i*_), to capture our knowledge of *q*_*i*_, with our best estimate of *q*_*i*_ being the expectation value ⟨*θ*_*i*_⟩.

The posterior distribution *P* (*θ*_*i*_|*D*) describes our knowledge of *θ*_*i*_, after seeing the data, *D*, i.e. the RNA-Seq read count data (*N, n*_*i*_),

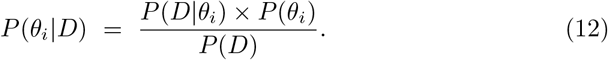

The posterior *P* (*θ*_*i*_|*D*) is determined by three factors: the prior probability given a hypothesis *H, P* (*θ*_*i*_|*H*), the likelihood of *θ*_*i*_, either denoted as *P* (*θ*_*i*_|*D*) or Λ(*θ*_*i*_), and a normalising factor *P* (*D*) in the denominator. This normalising factor is called the evidence or the marginal likelihood.

To find a likelihood function *P* (*D*|*θ*_*i*_) for all possible *θ*_*i*_, we can use the binomial distribution to model *P* (*D*|*θ*_*i*_) as a simple binomial process,

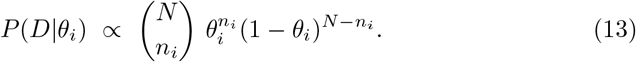

The prior distribution captures our knowledge about the system before we collect data. A convenient choice for the prior is the Beta distribution, as it is the conjugate distribution of the binomial distribution, owing to it having the same functional form. This allows us to proceed analytically. The prior distribution can be written as

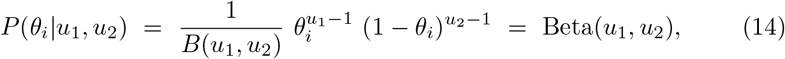

where the hyperparameters *u*_1_, *u*_2_ ∈ ℝ^+^, can be used to capture existing knowledge, if available. Here, we choose a bias-free, flat prior (*u*_1_ = *u*_2_ = 1).

Finally, the evidence *P* (*D*) is the probability that the data *D* is produced. It can be calculated by integrating over the numerator in equation 12,

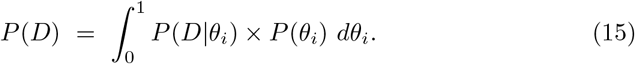

For two datasets, *D*_1_ and *D*_2_, we can find a posterior distribution *P* (*θ*_*i*_|*D*) for Hypothesis 1 and Hypothesis 2 separately. For Hypothesis 1, the assumption is that the same expression probability can explain both datasets, *θ*_*i*_ ⇝ *D*_1_ and *θ*_*i*_ ⇝ *D*_2_, resulting in

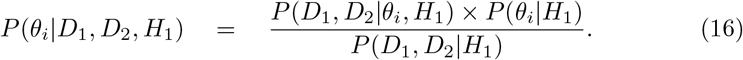

For Hypothesis 2, we assume that the expression probabilities are different between experiments, *θ*_*i*1_ ⇝ *D*_1_ and *θ*_*i*2_ ⇝ *D*_2_,

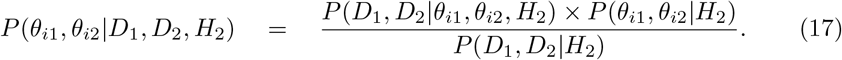

We define a prior for the single expression probability in Hypothesis 1,

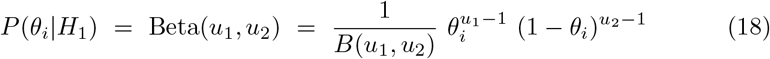

and the prior for two expression probabilities in Hypothesis 2,

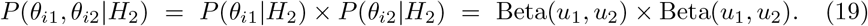

Note that we have assumed the same priors over *θ*_*i*1_ and *θ*_*i*2_. The likelihood for the data (*D*_1_, *D*_2_) given Hypothesis 1 can be written as

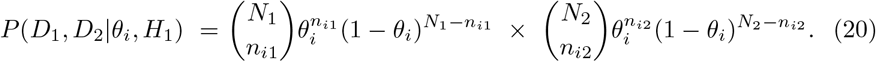

For Hypothesis 2, the likelihood of the data given *H*_2_ depends on the two expression parameters of the model, *θ*_*i*1_ and *θ*_*i*2_,

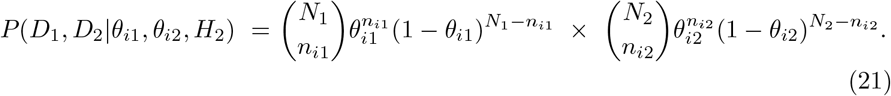

Thus, the posterior of Hypothesis 1 simplifies, thanks to conjugate priors, to the following equation,

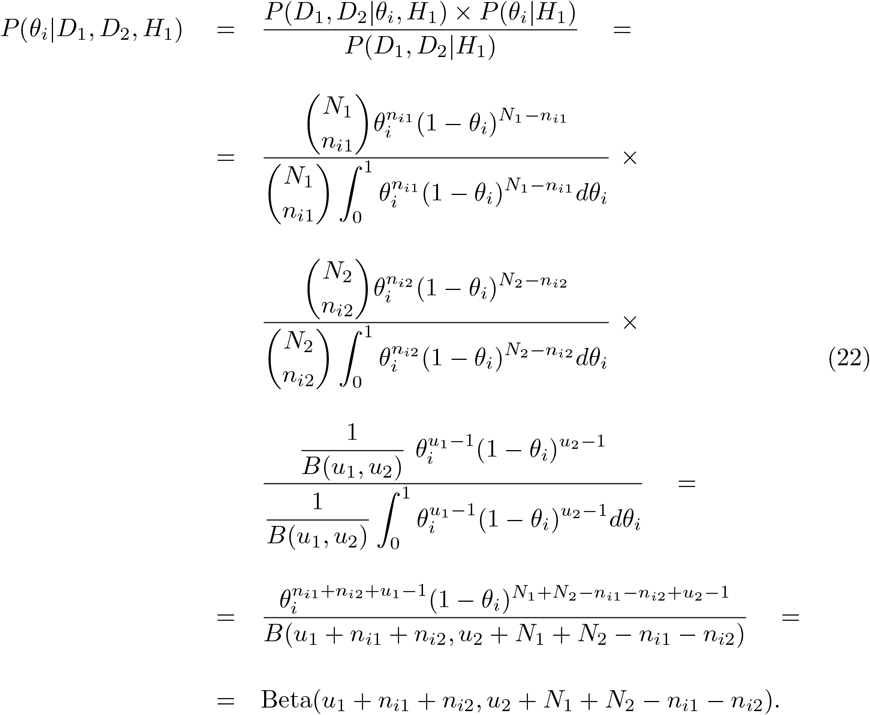

We used the integral definition of the Beta function to simplify the denominator. The posterior can now be expressed as a Beta distribution, compare Equation 14. Analogously, for Hypothesis 2 we can formulate and simplify the posterior probability distribution to

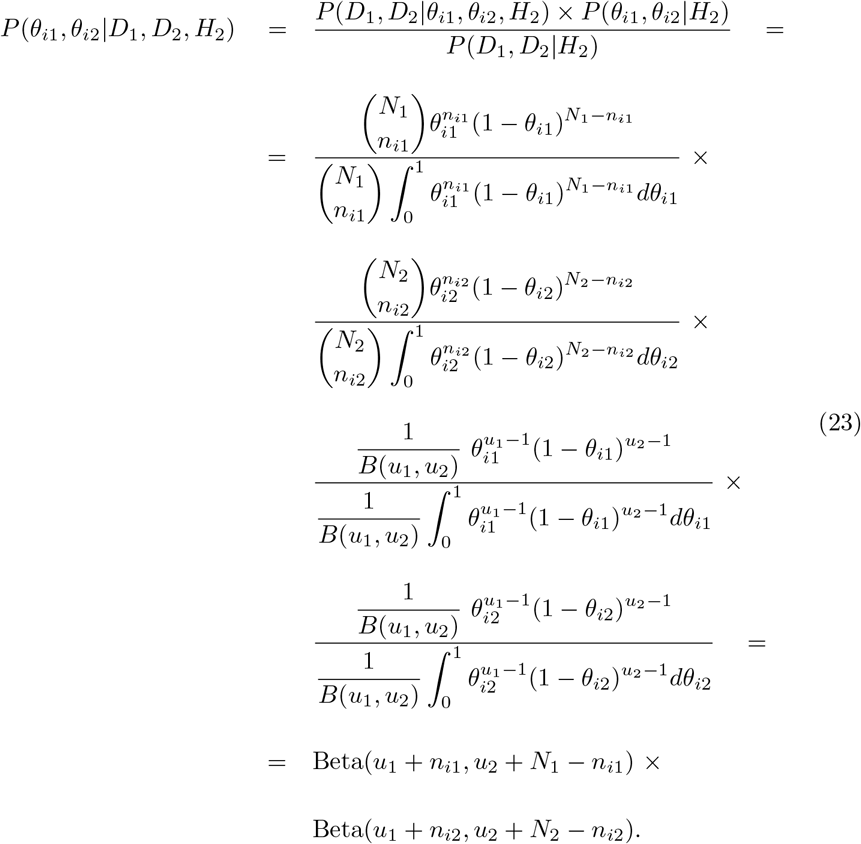

The Bayes factor (*BF*_21_) is a ratio of the statistical evidences *P* (*D*|*H*_1_) and *P* (*D*|*H*_2_) supporting the two hypotheses. We already found the analytical expressions for the evidences as the denominator of their posteriors.

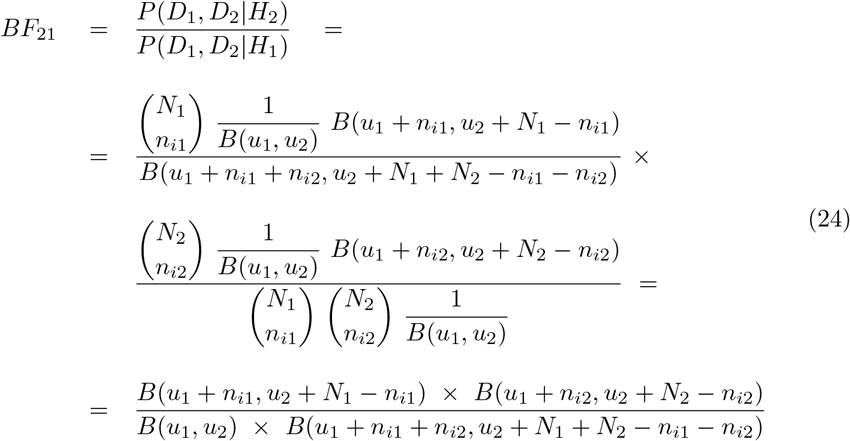

Where all except one pre-factor, *B*(*u*_1_, *u*_2_), cancel. For a flat prior, *B*(*u*_1_, *u*_2_) = 1.

As is common practice in differential gene expression analysis, we proceed to calculate an inferred log_2_ fold change (*iFC*). Using the same prior as above and integrating over *θ* to compute the expectation value, results in

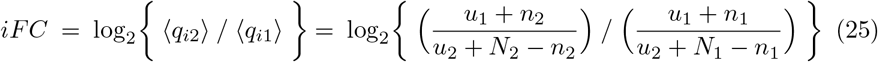

This approach allows for replicates to be handled by summing the *N* and *n*_*i*_ values. From these combined values, we can infer an average expression rate, log_2_ fold change (*iFC*) and compute a Bayes factor (*BF*_21_) between different conditions.

### Extension to multiple models to evaluate the consistency of expression across replicates

The framework described for two datasets can also be used on *k* datasets and with one against *k* number of models. This can be used, for example, how described in the Results section, to test the consistency of a gene’s expression across replicates in an RNA-Seq experiment.

We define two hypotheses. **Hypothesis H**_**1**_ states that the data from all replicates can be explained by a statistical model with only one expression probability for each gene,

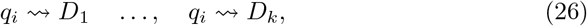

i.e the data are consistent with the biological and technical variance that might be expected between replicates. **Hypothesis H**_**k**_ states the data are best explained by a separate expression probability for each replicate,

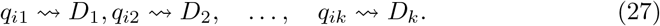

In principle, the number of models need not be equal to the number of replicates, and any number greater than 1 and less or equal to *k* could be explored, i.e. we would be asking whether various subsets of replicates are consistent. Due to the combinatorics for large numbers of replicates (see below), we limit ourselves here to the extreme case of every replicate being different. If the replicates are consistent, we can simply sum all the *n*_*ir*_ for replicate *r* for each gene *i*, and likewise for the *N*_*r*_ with the above framework.

For each gene we calculate a Bayes factor, describing how much the RNA-Seq data supports Hypothesis *H*_*k*_ over Hypothesis *H*_1_, i.e. how consistent the expression is between replicates.

For **Hypothesis H**_**1**_ the evidence is given by

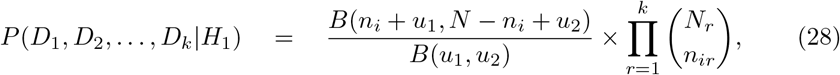

with *N* = _*r*_ *N*_*r*_ and *n*_*i*_ = _*r*_ *n*_*ir*_ for all replicates *r*. The evidence for **Hypothesis H**_**k**_ is given by

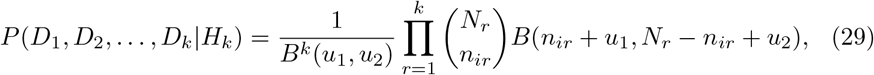

with *N* = ∑ _*r*_ *N*_*r*_ and *n*_*i*_ = ∑ _*r*_ *n*_*ir*_. By computing the ratio of the evidence for Hypothesis *H*_*k*_ over Hypothesis *H* _*k*_, we have a general way of testing *k* models over 1 model,

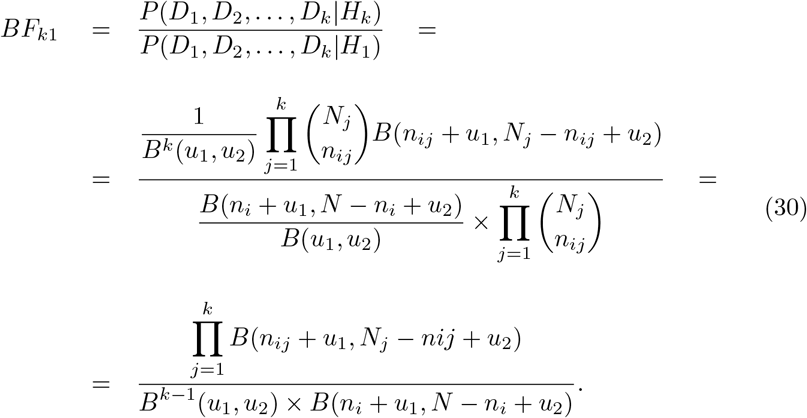

### 5.2 Bayes factors do not require a correction for the length of genes

In Section 2.4 we stated that Bayes factors grow with the evidence given in data to calculate them. The more reads map to a gene, for example, the stronger the evidence we can find to support the hypotheses. Therefore, we investigated the relationships between gene lengths, q-values and Bayes factors, because longer genes have a higher probability of reads mapping to them. We found, however, that in practice the analysis is not sensitive to gene length and no further normalisation is required, Supplementary Figure 9. We document an extreme example of what happens if the total read depth between conditions or treatments varies and how this is not an issue for the framework in Figures 6 and 7.

**Figure 9:**
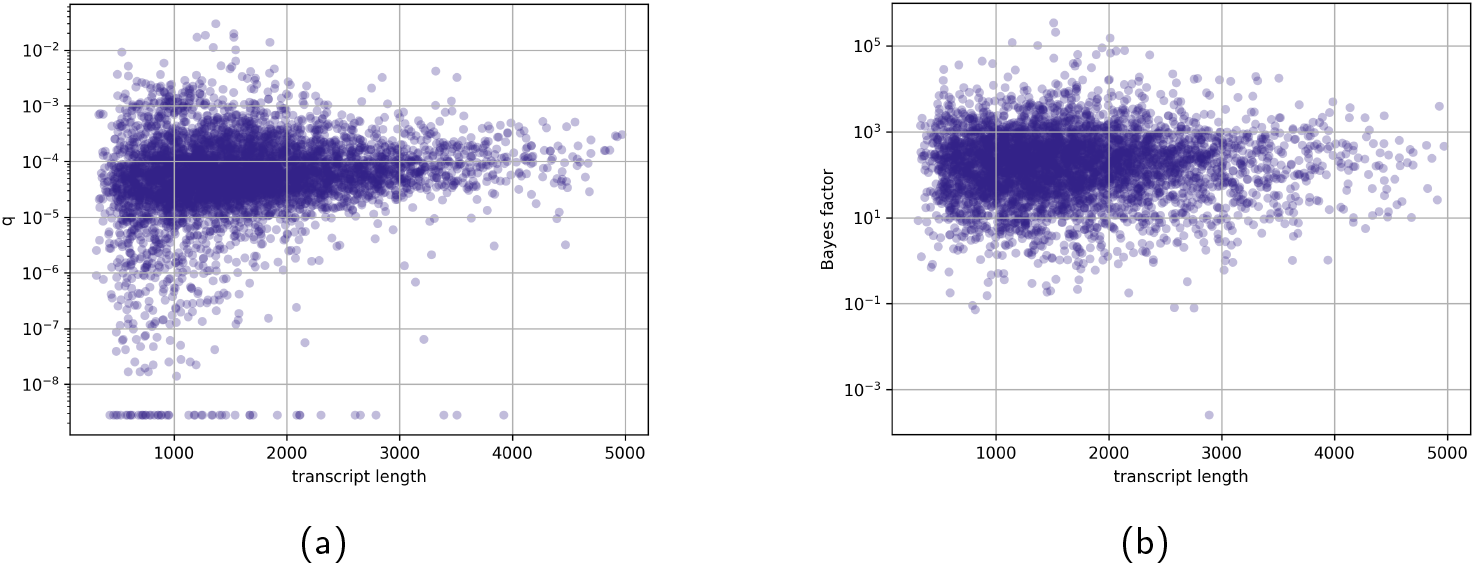
Bayes factors for differential expression (*BF*_21_) do not require gene length corrections. (a) We did not find a correlation between transcript lengths and expresson probability, *q*, of genes (log-scale, across all replicates in the Yeast dataset [15]) and neither can we find a correlation in (b) between transcript lengths and Bayes factors of genes (log-scale, across all replicates). Shown are 5219 of the 7126 yeast genes. We conclude that Bayes factors are not gene length dependent and that no additional normalisation is necessary.

## Supplementary Material

## Acknowledgements

We thank Hugh Woolfenden for the R implementation. We thank our colleagues Ander Movilla-Miangolarra and Pirita Paajanen for their comments and critical discussions on the manuscript. Furthermore, we would like to mention Emma Raven, Marina Mill’an-Blanquez, Burkhard Steuernagel, Philippa Borrill, Marek Glombik, Christine Faulkner and Hugh Woolfenden for many insightful conversations on the topic over the last years. Finally, we thank the JIC and TSL community – above all, the members of the Morris Lab – for thought-provoking discussions that shaped this work.

